# Istradefylline, an adenosine A2a receptor antagonist, ameliorates neutrophilic airway inflammation and psoriasis in mice

**DOI:** 10.1101/2021.05.21.445220

**Authors:** Mieko Tokano, Masaaki Kawano, Rie Takagi, Sho Matsushita

## Abstract

**Objective:** Extracellular adenosine is produced from secreted ATP by cluster of differentiation (CD)39 and CD73. Both are critical nucleotide metabolizing enzymes of the adenosine generating pathway and are secreted by neuronal or immune cells. Adenosine plays a role in energy processes, neurotransmission, and endogenous regulation of inflammatory responses. Istradefylline is a selective adenosine A2a receptor (A2aR) antagonist used for the treatment of Parkinson’s disease. We have reported that adenosine primes hypersecretion of interleukin (IL)-17A via A2aR. Istradefylline, as well as an inhibitor of CD39 (ARL67156) and an inhibitor of CD73 (AMP-CP), suppressed IL-17A production, and the administration of istradefylline to mice with experimental autoimmune encephalomyelitis (EAE) led to the marked amelioration of the disease. These previous results suggest that adenosine is an endogenous modulator of neutrophilic inflammation. We investigated the effect of istradefylline, ARL67156 and AMP-CP on other mouse models of neutrophilic inflammation.

**Methods:** We tested the effect of istradefylline, ARL67156 and AMP-CP on OVA-induced neutrophilic airway inflammation or imiquimod (IMQ)-induced psoriasis in mice. These two model mice received these drugs orally or percutaneously, respectively. The production of IL-17A in the lung and ear thickness were used as an index of the effects.

**Results:** We show that istradefylline, ARL67156 and AMP-CP suppressed the OVA-induced IL-17A production in the lung and IMQ-induced psoriasis.

**Conclusion:** These results indicate that adenosine-mediated IL-17A production plays a role in neutrophilic inflammation models, and moreover, istradefylline, ARL67156, and AMP-CP are effective in animal models of neutrophilic inflammation. Some clinical relevancies in COVID-19 are discussed.

## Introduction

Adenosine, a molecular moiety of ATP, ADP, and AMP, is involved in energy processes and is essential for the phenomena of life. Extracellular adenosine is produced from secreted ATP by ectonucleotidases, such as ecto-nucleoside triphosphate diphosphohydrolase (E-NTPDase)-1 cluster of differentiation (CD)39, which converts ATP or ADP to ADP or AMP, respectively, and the 5’-nucleotidase CD73, which dephosphorylates AMP to adenosine. CD39 and CD73 are expressed on the surface of endothelial cells^1,2)^ and immune cells^3–5)^. Adenosine binds to adenosine receptors expressed on the cell surface. There are four subtypes of adenosine receptors, A1, A2a, A2b, and A3, which belong to a superfamily of membrane proteins called the G protein-coupled receptor family. A2aR and A2bR signal the Gs protein to trigger cAMP synthesis. On the other hand, A1R and A3R signal the Gi protein to trigger cAMP degradation^6)^. A1R, A2bR and A3R are widely expressed in the body. In contrast, A2aR is expressed at high levels in only a few regions of the body, namely the striatum, olfactory tubercle, nucleus accumbens, endothelial cells, vascular smooth muscle cells, platelets, and immune cells^7)^. A1R and A2aR are high-affinity receptors, whereas A2bR and A3R are low-affinity receptors^8,9)^.

The purine nucleoside adenosine also plays a role as a neurotransmitter, primarily in the striatum, olfactory tubercle and nucleus accumbens^10)^. Istradefylline is a selective A2aR antagonist used for the treatment of Parkinson’s disease^11)^. Furthermore, adenosine is a potent endogenous regulator of inflammation and immune reactions^6)^. However, the molecular mechanisms underlying its effects are largely unknown. In previous a study, adenosine was reported to induce T-helper (Th)17 differentiation by activating A2bR^12)^.

Th17 cells are a subset of T-helper cells that differentiate from naïve CD4^+^ T cells in the presence of tumor growth factor (TGF)-β and interleukin (IL)-6. These cytokines are secreted by antigen-presenting cells in response to stimulation via T cell receptor (TCR) antigen^13–15)^. IL-17A production by Th17 cells drives neutrophil recruitment and neutrophilic inflammation^16,17)^. The IL-17A-mediated responses are induced in receptor-expressing cells, such as endothelial cells, epithelial cells, and fibroblasts^18)^. Neutrophilic inflammation is associated with many diseases^19)^, including autoimmune diseases^20–23)^, neutrophilic airway inflammation^24,25)^, psoriasis^26,27)^, severe atopic dermatitis^28)^, and multiple sclerosis^29–34)^. There are currently no specific therapies that use low-molecular weight chemicals for neutrophilic inflammation, nevertheless corticosteroids are a specific therapy for eosinophilic inflammation. However, recent studies by ourselves and others suggested that dopamine D1-like receptor antagonists and dopamine D2-like receptor agonists suppress neutrophilic inflammation by suppressing Th17 differentiation and activation^35–37)^. We recently reported that adenosine is also produced by activated CD4^+^ T cells, mainly during T cell-APC interactions, primes the hypersecretion of IL-17A by CD4^+^ T cells, where A2aR plays a role in the hypersecretion of IL-17A. Istradefylline, an inhibitor of CD39 (ARL67156), and an inhibitor of CD73 (AMP-CP) suppressed IL-17A production, and the administration of istradefylline to mice with experimental autoimmune encephalomyelitis led to the marked amelioration of symptoms^38)^. These results suggest that adenosine is an endogenous modulator of neutrophilic inflammation.

In this study, we tested the effect of istradefylline, ARL67156, and AMP-CP on other models of neutrophilic inflammation, such as OVA-induced neutrophilic airway inflammation and imiquimod-induced psoriasis. We show that istradefylline, ARL67156 and AMP-CP are effective in animal models of neutrophilic inflammation.

## Materials and methods

### Mice

OVA TCR-transgenic DO11.10 mice were obtained from The Jackson Laboratory (Bar Harbor, ME). C57BL/6 mice were obtained from Japan SLC (Shizuoka, Japan). Mice were housed in appropriate animal care facilities at Saitama Medical University and handled according to the international guidelines for experiments with animals. All experiments were approved by the Animal Research Committee of Saitama Medical University.

### Measurement of cytokine concentrations in the lung

Airway inflammation was induced as described previously^36)^. Briefly, eight-week-old female DO11.10 mice received a subcutaneous inguinal injection (100 μg/mouse) of 2 mg/mL OVA (Sigma) in PBS (-) emulsified in complete Freund’s adjuvant (CFA) containing mycobacterium tuberculosis H37Ra (100 μg/mouse; Difco) on day −8. Mice also received oral PBS (-), an A2aR antagonist (Istradefylline) (6 μg/mouse), an inhibitor of CD39 (ARL67156, Tocris) (0.5 mg/mouse) or an inhibitor of CD73 inhibitor (adenosine 5’-(α, β-methylene) diphosphate (AMP-CP; Tocris) (0.5 mg/mouse) on days −10, −8, −6, −4, −2, and −1. Mice were challenged with an aerosolized solution of 3% OVA or PBS (-) for 10 min on day −1. The mice were analyzed on day 0. Lung cells were prepared as previously described^36)^. Briefly, the left lungs were cut out, homogenized, and incubated in 10 mL of DMEM medium containing 10% FCS, 100 U/mL penicillin, 100 μg/mL streptomycin, 1 mM sodium pyruvate, 50 μM 2-mercaptoethanol, 50 μg/mL gentamycin, 1 μg/mL amphotericin, and collagenase from clostridium histolyticum (Sigma-Aldrich) for one hour. Following incubation, the lung lymphocytes were washed twice. Lung lymphocytes (1 × 10^6)^ were seeded in a round-bottomed 96-well plate and then incubated in in 500 μL of DMEM medium containing 10% FCS, 100 U/mL penicillin, 100 μg/mL streptomycin, 1 mM sodium pyruvate, 50 μM 2-mercaptoethanol, 50 μg/mL gentamycin, and 1 μg/mL amphotericin for four days. The supernatant fluids obtained by lung homogenates were then collected for the IL-17A, IFN-γ, and IL-5 ELISAs.

### Histological examination

The histological examination was performed as previously reported^36)^. The right lungs were resected, fixed with 10% neutralized buffered formalin (Wako), and embedded in paraffin. Three-micrometer-thick sections were stained with hematoxylin and eosin. The numbers of polymorphonuclear leukocytes (other than eosinophils) per 2500 μm^2^ were counted.

### The mouse model of imiquimod (IMQ)-induced psoriasis

Psoriasis was induced in the mouse model as previously described^39)^. Briefly, C57BL/6 mice were treated with either IMQ cream containing 5% IMQ (Mochida Pharmaceutical) or sham cream, which was applied on the ears for 5 consecutive days. On day 9, the ear thickness (μm) was measured. In the treatment groups, a cream containing 5% A2aR antagonist (Istradefylline), liquid containing 10 mM CD39 inhibitor (ARL67156), or liquid containing 10 mM CD73 inhibitor (AMP-CP) was used. The histological examination was performed as previously reported^36)^. On day 9, the ears were resected, fixed with 10% buffered and neutralized formalin (Wako), and embedded in paraffin. Three-micrometer-thick sections were stained with hematoxylin and eosin.

### Cytokine ELISAs

The concentrations of IFN-γ, IL-5, and IL-17A in cell supernatants were measured using specific ELISA kits (DuoSet Kit, R&D). Any value below the lower limit of detection (15.6 pg/mL) was set to 0. No cytokine cross-reactivity was observed within the detection ranges of the kits. If necessary, samples were diluted appropriately so that the measurements fell within the appropriate detection range for each cytokine.

### Statistical analysis

Differences between two groups were analyzed using an unpaired Student’s *t*-test. Differences between three or more groups were analyzed using a one-way ANOVA with Tukey’s post-hoc test. Clinical scores were analyzed using a non-parametric Mann-Whitney U-test. All calculations were performed using KaleidaGraph software program (Synergy software, Reading, PA, USA). P values of <0.05 were considered to indicate statistical significance.

## Results

### An adenosine A2a receptor antagonist, istradefylline, suppresses OVA-induced neutrophilic airway inflammation in DO11.10 mice

First, we tested the effect of an adenosine A2a receptor antagonist, istradefylline, on OVA-induced neutrophilic airway inflammation in OVA TCR-transgenic DO11.10 mice. DO11.10 mice were challenged with nebulized OVA or with PBS as a control. The administration of istradefylline was performed starting from 10 days before nebulization (Fig.1a). Our previous study showed a clear correlation between IL-17A in the lung and neutrophilic airway inflammation^36)^. Indeed, the concentration of IL-17A increased in the lungs of OVA-challenged DO11.10 mice, which were suppressed by istradefylline on day 4 (Fig.1b). Time course studies showed that the production of IL-17A was time-dependent (Fig.1c). We observed that istradefylline treatment suppressed IL-17A (a Th17-related cytokine) and IFN-γ (a Th1-related cytokine) secretion on day 4 and had no significant effect on IL-5 (a Th2-related cytokine) secretion (Fig.1d).

**Figure 1.**
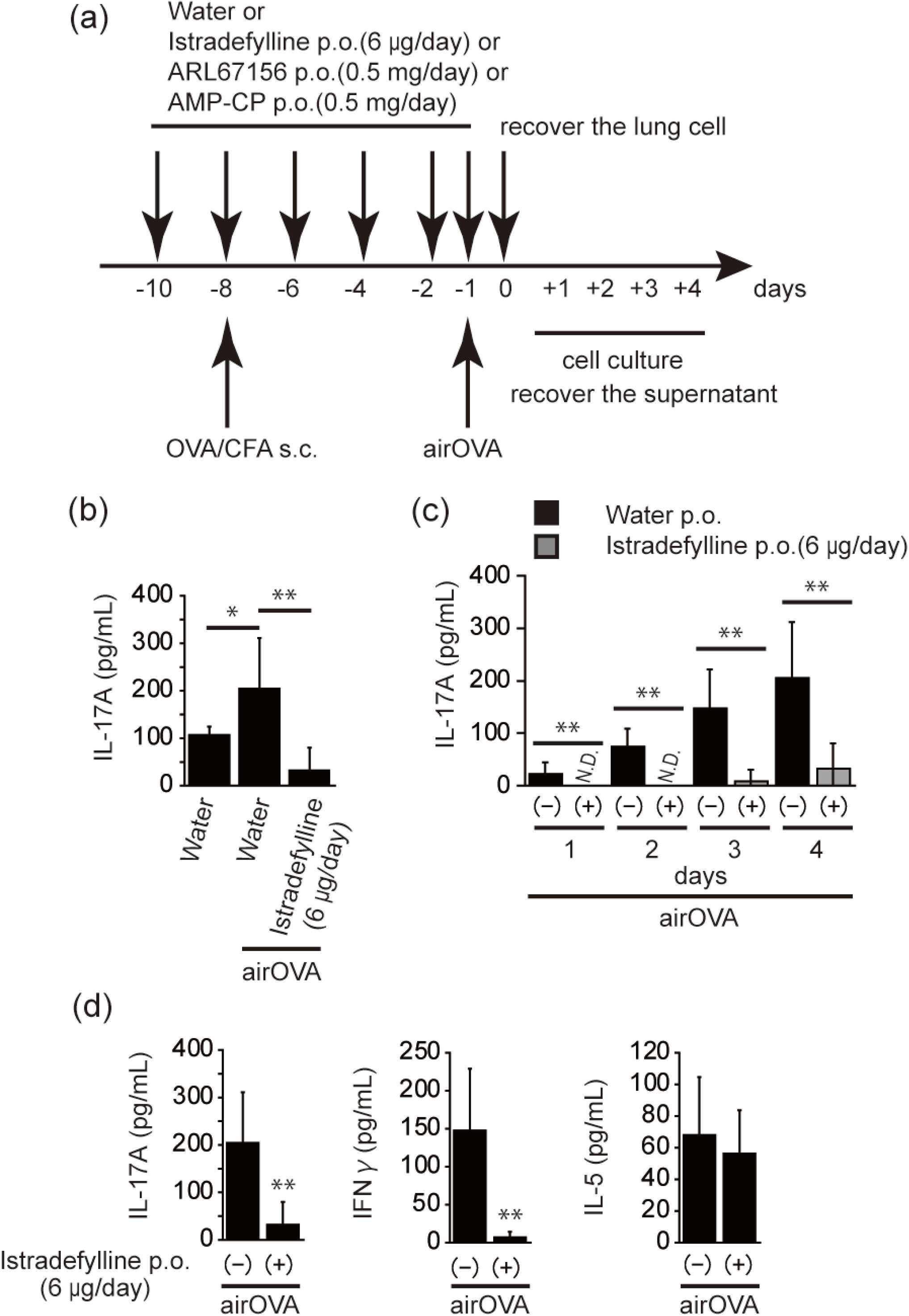
An adenosine A2a receptor antagonist, istradefylline, suppresses OVA-induced neutrophilic airway inflammation in DO11.10 mice. (a) The protocol of the OVA-induced neutrophilic airway inflammation assay. (b, c) Lung homogenate was assayed for concentrations of IL-17A by an ELISA. (d) Lung homogenate was assayed for concentrations of IL-17A (left), IFN-γ (center), or IL-5 (right) by an ELISA. The experiments were repeated 7-10 times, and similar results were obtained. Data are expressed as the mean ± SD and were compared using an unpaired Student’s *t*-test. \**p* < 0.05 and \*\**p* < 0.01, in comparison to the value of water (challenged OVA).

### Istradefylline suppresses OVA-induced neutrophil infiltration in DO11.10 mice

The histology of OVA-challenged DO11.10 mice showed prominent neutrophil infiltration into the peribronchial area (Fig. 2a), while the infiltration declined in mice that received istradefylline (Fig. 2b, 2c; *p*=0.021). Accordingly, istradefylline-treatment suppressed neutrophilic airway inflammation.

**Figure 2.**
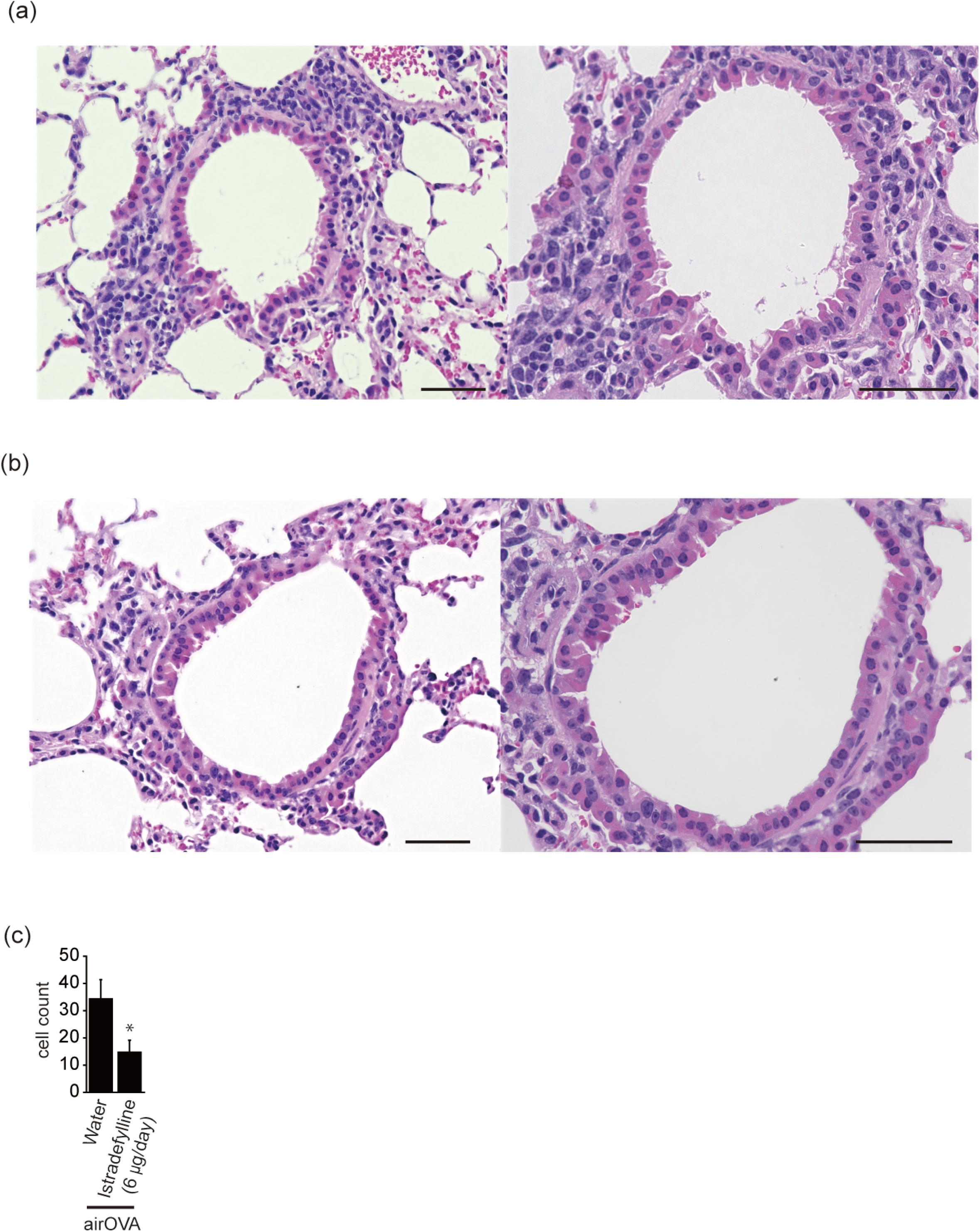
Istradefylline suppresses OVA-induced neutrophilic infiltration in DO11.10 mice. The lung sections from mice administered oral water (a) or istradefylline solution (b) were stained with hematoxylin and eosin (Scale bar, 50 μm). (c) The numbers of polymorphonuclear leukocytes per 2500 μm^2^. The experiments were repeated three times, and similar results were obtained. Data are expressed as the mean ± SD and were compared using an unpaired Student’s *t*-test. \**p* < 0.05, in comparison to the value of water (challenged OVA).

### ARL67156 and AMP-CP also suppress OVA-induced neutrophilic airway inflammation in DO11.10 mice

Since we found that istradefylline suppressed the production of IL-17A in the lung, we next examined the effect of a CD39 inhibitor (ARL67156) and a CD73 inhibitor (AMP-CP) on OVA-induced neutrophilic airway inflammation. ARL67156 and AMP-CP inhibit the production of adenosine (data not shown). DO11.10 mice were challenged with nebulized OVA, and the administration of ARL67156 and AMP-CP was performed from 10 days before OVA nebulization. As in the case of istradefylline treatment, ARL67156 and AMP-CP treatment suppressed the production of IL-17A in the lung (Fig.3). This suggests that adenosine promotes neutrophilic airway inflammation by hypersecretion of IL-17A.

**Figure 3.**
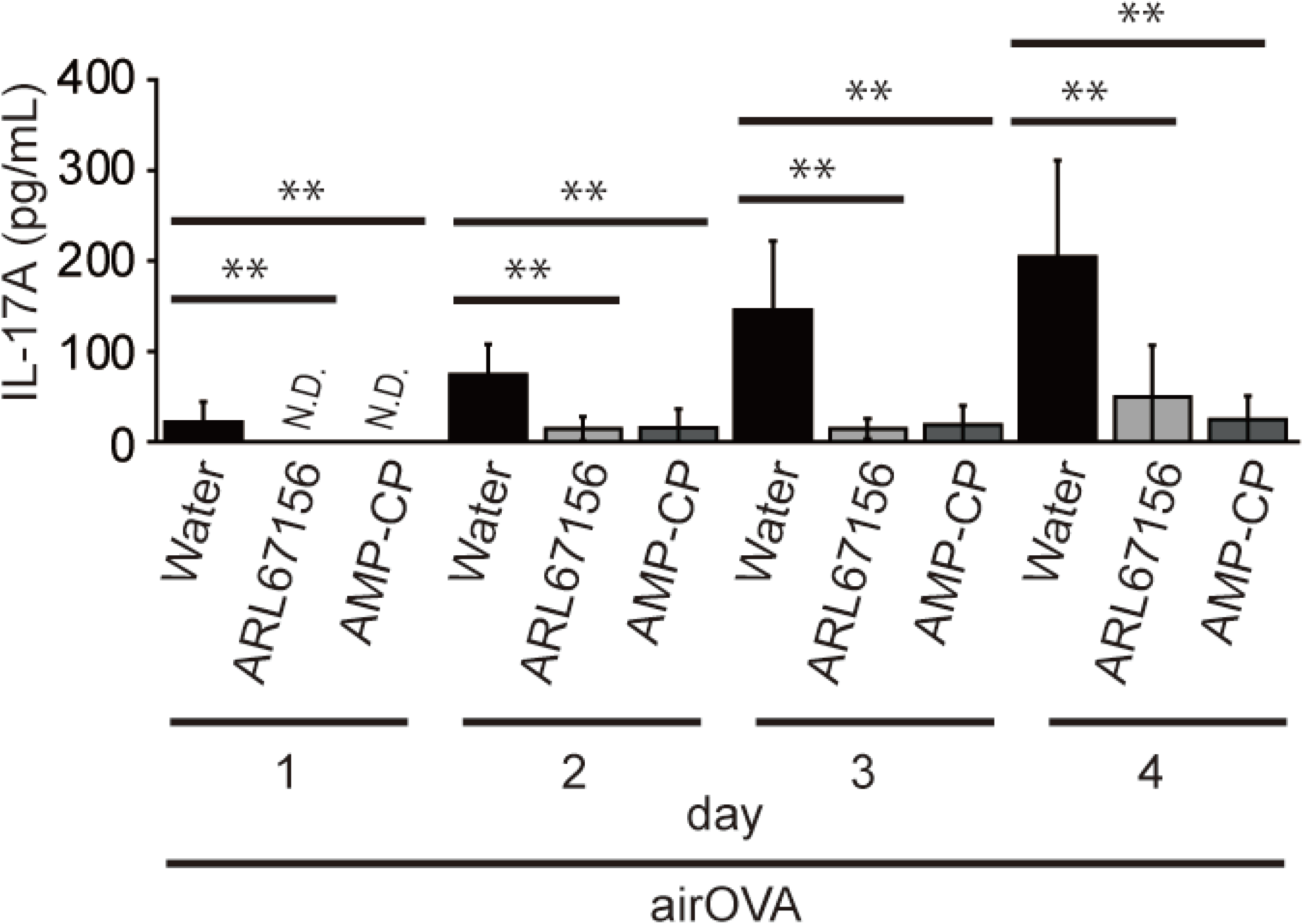
Inhibitor of CD39 (ARL67156) and inhibitor of CD73 (AMP-CP) suppresses OVA-induced neutrophilic airway inflammation in DO11.10 mice. DO11.10 mice were treated as described for Fig.1. Lung homogenate was assayed for concentrations of IL-17A by an ELISA. The experiments were repeated 7-10 times, and similar results were obtained. Data are expressed as the mean ± SD and were compared using an unpaired Student’s *t*-test. *P < 0.05 and **P < 0.01 in comparison to the value of water (challenged OVA).

### Istradefylline, ARL67156, and AMP-CP suppress imiquimod (IMQ)-induced psoriasis in mice

Psoriasis is a Th17-mediated disease^26,27)^. Indeed, the skin infiltration of neutrophils, activated monocytes, Th17 cells are observed in psoriasis and a mouse model of IMQ-induced psoriasis ^40–42)^. Mice were treated with either 5% IMQ cream or sham cream. Neutrophilic inflammation and hyperkeratosis of the skin were induced by IMQ (Fig. 4a, b, c). In the treatment groups, 5% istradefylline-containing cream, 10 mM ARL67156 or 10 mM AMP-CP-containing liquid was used. Istradefylline, ARL67156, and AMP-CP significantly suppressed the effect of IMQ (Fig. 4d). All these observations collectively suggest that the oral or transdermal administration of istradefylline, ARL67156, and AMP-CP suppresses Th17-mediated disease.

**Figure 4.**
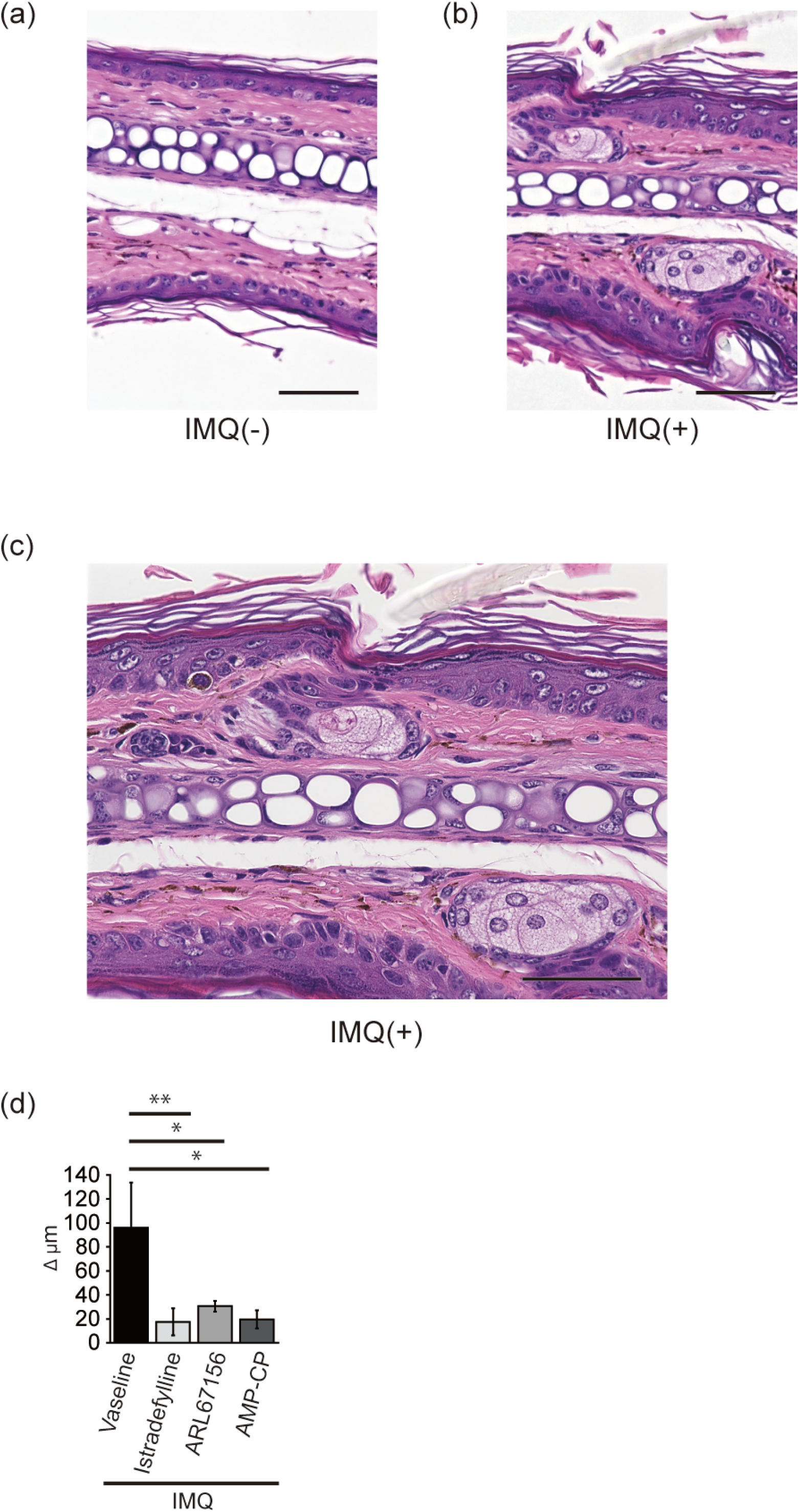
Istradefylline, ARL67156, and AMP-CP suppress IMQ-induced psoriasis in mice. Mice were treated either with sham cream (a) or IMQ cream containing 5% IMQ (b, c). Figure 4c shows a close-up view of Figure 4b. Ear sections from mice were stained with hematoxylin and eosin (Scale bar, 50 μm). (d) The ear thickness (μm) of was measured on day 9. Values obtained by subtracting the negative control values (Δμm) are shown. In treatment groups, cream containing 5% istradefylline, liquid containing 10 mM ARL67156, or liquid containing 10 mM AMP-CP inhibitor was used. Data were obtained from three independent experiments (n = 3-4 mice/group), and similar results were obtained. Data are expressed as the mean ± SD and were compared using an unpaired Student’s *t*-test. *P < 0.05 and **P < 0.01, in comparison to the value of the non-treatment group.

## Discussion

Atopic asthma is usually triggered by allergens or by antigen-non-specific stimuli, in which Th2 inflammation, group 2 innate lymphoid cell (ILC2) activation and eosinophilic inflammation play a pivotal role. Approximately 50% of elderly and 90% of young individuals with asthma show the atopic phenotype. On the other hand, the recruitment and activation of neutrophils in airways are associated with resistance to corticosteroids. Approximately 40% of elderly patients with asthma have neutrophilic airway inflammation^24,25,43,44)^, accompanying increased bronchial IL-17^+^ cells^45–47)^. The TCR-transgenic DO11.10 mice have TCR, which specifically recognizes MHC class II-OVA peptide complex. OVA nebulization alone could induce IL-17-dependent neutrophilic airway inflammation^28,36,48–50)^. This response is OVA-specific, as other antigens could not induce neutrophilic airway inflammation. In addition, deletion of the IL-17 gene suppressed the neutrophilic airway inflammation^50)^. Thus, this animal model is similar to the pathogenesis of antigen-induced Th17-mediated neutrophilic airway inflammation^36)^. Our studies demonstrate that istradefylline-treatment suppressed IL-17-dependent neutrophilic airway inflammation in DO11.10 mice. Similarly, ARL67156 and AMP-CP, which inhibit the production of adenosine, suppressed IL-17-dependent neutrophilic airway inflammation, which corroborates our previous findings^38)^. Furthermore, the modulation of signaling via A2aR might ameliorate autoimmune diseases, including allergy and infections. The latter may include disseminated intravascular coagulation (DIC) or acute respiratory distress syndrome (ARDS) in SARS-CoV-2 disease (COVID-19), which is reportedly associated with neutrophil extracellular traps (NETs)^51–54)^. In recent previous studies, patients with severe COVID-19 showed the aberrant activation of neutrophils and Th17 promotion^55)^, and IL-17 can serve as a biomarker of the severity of COVID-19^56)^. Indeed, autopsy samples from the lungs of COVID-19 patients showed neutrophil infiltration in pulmonary capillaries^57)^, and the peripheral blood of patients showed an increased frequency of Th17 cells^58)^. Accordingly, it is conceivable that istradefylline-treatment may suppress IL-17 secretion and neutrophilic airway inflammation in COVID-19.

Psoriasis had long been characterized as a Th1-mediated disease because psoriatic lesions showed the elevated mRNA expression of Th1 cytokines (IFN-γ and TNF-α)^59)^. Recent studies have shown that the pathology of psoriasis is strongly dependent on IL-17A^60)^. In an IMQ-induced mouse model, activated Th17 cells and marked skin infiltration of neutrophils are observed^40,42)^. Our studies demonstrate that istradefylline, ARL67156, and AMP-CP suppress IMQ-induced murine psoriasis. It is therefore conceivable that adenosine promotes IL-17A production in an IMQ-induced mouse model. We also confirmed that γδT cells secreted IL-17A after stimulation with agonistic anti-CD3/CD28 antibodies in the presence of adenosine (data not shown). In the dermis with psoriasis, IL-23 from keratinocytes, activated Langerhans cells, macrophages, and dendritic cells are capable of promoting the production of IL-17A by γδT cells^61–63)^. Adenosine-mediated IL-17A production may play an important role in psoriasis.

Our study demonstrated that istradefylline as well as ARL67156 and AMP-CP suppress neutrophilic airway inflammation and psoriasis in mice, which strongly attests to the *in vivo* relevance of adenosine-mediated IL-17A production. It is also suggested that istradefylline as well as ARL67156 and AMP-CP may be effective treatments for Th17-mediated diseases, such as psoriasis, neutrophilic bronchial asthma, and autoimmune diseases, due to their suppression of the hypersecretion of IL-17A from Th17 cells. Furthermore, we reported that adenosine is produced by activated CD4^+^ T cells and primes hypersecretion of IL-17A by CD4^+^ T cells via A2aR^38)^. These results suggest that CD39 and CD73 expressed on the CD4^+^ T cell surface converts ATP to adenosine, and adenosine binds to A2aR and primes hypersecretion of IL-17A (Fig. 5). Some researchers argue that an A2aR agonist, CGS21680, suppresses Th17 differentiation^64–66)^. Because CGS21680 is much less selective than the A2aR agonist we used in a recent previous study (PSB0777)^38)^, it is highly conceivable that these studies gave contradictory results. In addition, some researchers argue that methotrexate exerts an anti-rheumatic effect by promoting adenosine release^6)^. Indeed, different expression patterns of dopamine receptor subtypes are observed on different populations of immune cells, depending on the activation status of cells^67)^. Because A1R agonism and A2aR antagonism are biologically equivalent in the presence of adenosine, it is conceivable that adenosine exhibits anti-rheumatic effects, provided A1R is dominantly expressed by T cells in the local environment of the rheumatic synovium. The expression patterns of adenosine receptor subtypes in our current animal models are under investigation.

**Figure 5.**
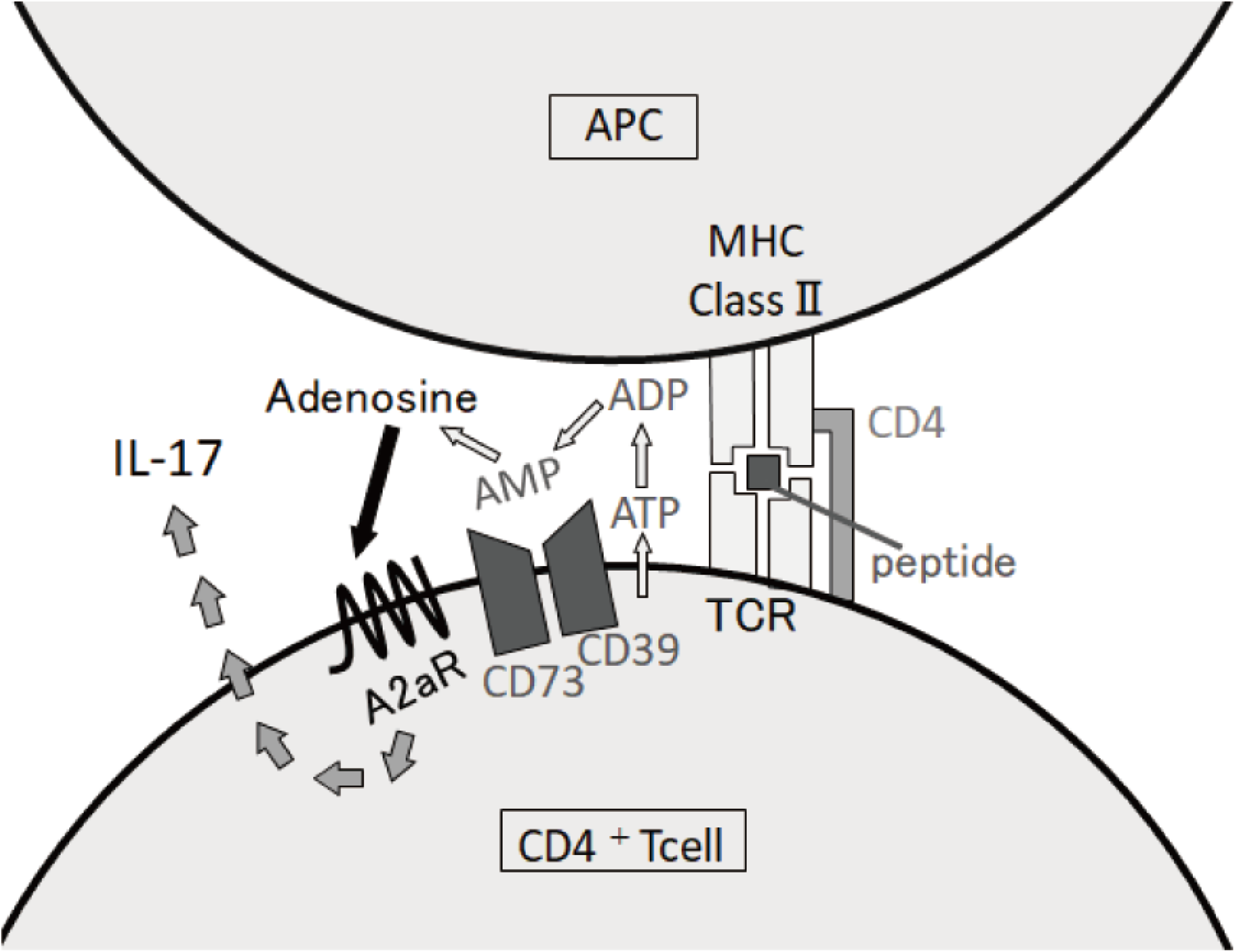
CD4^+^ T cells produce ATP by antigen presentation. Extracellular adenosine is produced from ATP secreted by CD39 and CD73. Adenosine binds to A2aR expressed on the cell surface and primes hypersecretion of IL-17A (hypothesis).

It is suggested that the concentrations of adenosine are increased as much as 50 times by physiological stimuli such as hypoxia, hypoglycemia, and ischemia^68)^. A previous study also suggested that extracellular adenosine is transported into the cell by transporters or that it is rapidly broken down by adenosine deaminase or adenosine kinase^69)^. It is probable that adenosine induces neutrophilic inflammation in acute stages, *i.e*., in the innate immunity-acquired immunity interface.

## Disclosure of ethical statements

No human participant was involved in this study.

## Acknowledgments

This work was supported by a Grant-in-Aid for Scientific Research (C) (no. 19K07201), awarded to M.K., a Grant-in-Aid for Young Scientists (B) (no. 18K15327) to R.T., and a Grant-in-Aid for Scientific Research (C) (no. 19K08887) awarded to S.M. by the Japanese Society for the Promotion of Science. This work was also supported by the 44th and 45th Science Research Promotion Fund, awarded to M.K. by the Promotion and Mutual Aid Corporation for Private Schools of Japan.

## Abbreviations

Th: T-helper
CD: cluster of differentiation
TGF: tumor growth factor
IL: interleukin
APCs: antigen presenting cells
MHC: major histocompatibility complex
TCR: T cell receptor
EAE: experimental autoimmune encephalomyelitis
CFA: complete Freund’s adjuvant
OVA: ovalbumin
IMQ: imiquimod
n: number of repeat experiments
SD: standard deviation.

## Author contributions

M.T., R.T., M.K., and S.M., performed the experiments. M.T., M.K., and S.M., conceived and designed the experiments. M.T., M.K., and S.M., wrote the manuscript. All authors discussed the results and commented on the manuscript.

